# Insights from single-strain and mixed culture experiments on the effects of heatwaves on freshwater flagellates

**DOI:** 10.1101/2024.06.14.598979

**Authors:** Lisa Boden, Chantal Klagus, Jens Boenigk

**Affiliations:** Department Biodiversity, University of Duisburg–Essen Universitätsstr. 5, 45141 Essen, Germany; Center for Water and Environmental Research, University of Duisburg–Essen Universitätsstr. 5, 45141 Essen, Germany

## Abstract

The increasing frequency and intensity of heatwaves driven by climate change significantly impact microbial communities in freshwater habitats, particularly eukaryotic microorganisms. Heterotrophic nanoflagellates are important bacterivorous grazers and play a crucial role in aquatic food webs, influencing the morphological and taxonomic structure of bacterial communities. This study investigates the responses of three flagellate taxa to heatwave conditions through single-strain and mixed culture experiments, highlighting the impact of both biotic and abiotic factors on functional redundancy between morphologically similar protist species under thermal stress. Our results indicate that temperature can significantly impact growth and community composition. However, density-dependent factors also had a significant impact. In sum, stabilizing effects due to functional redundancy may be pronounced as long as density-dependent factors play a minor role and can be overshadowed when flagellate abundances increase.

## Introduction

Microbial communities frequently experience and cope with natural fluctuations in abiotic environmental conditions (Ruokolainen *et al*. 2009). However, strong changes caused by climate change and anthropogenic activities put ecosystems under severe pressure (IPBES 2019). Microorganisms have evolved mechanisms to withstand changing environmental conditions (Bernhardt *et al*. 2020), but tolerances to abiotic stressor vary strongly (Díaz-Almeyda *et al*. 2017; Mascarin *et al*. 2018; Jain and Saraf 2021). Consequently, abiotic disturbances can result in lower abundance of individual species in the microbial community or total loss of species (Schulhof *et al*. 2020; Proesmans *et al* 2023). Such changes in biodiversity can significantly impact ecosystem functions and food webs (Mooney *et al*. 2009; IPBES 2019). However, the impact of losing individual species can be lowered by the presence of other species in the community that perform the same function, an attribute known as functional redundancy (Walker 1992, Naeem 1998). Functional redundancy is hypothesized to enhance the resilience and stability of ecosystems during the occurrence of stressors by maintaining ecological functions despite changes in taxon composition (Briggs *et al*. 2020, Chen *et al*. 2022).

As a major abiotic stressor, increased temperatures threaten biodiversity across different habitats (Urrutia-Cordero *et al*. 2017; Maire *et al*. 2022; Lau *et al*. 2024). In addition to the long-term increase of the average global surface temperature, more frequent and intense heatwaves have been predicted as a consequence of climate change (Woolwa *et al*. 2021; IPCC 2021). Heatwaves are short but intense increases of temperature and represent one of the most important challenges in aquatic habitats (Sun and Arnott 2022). During the occurrence of a heatwave, organisms are rapidly pushed beyond the limit of their temperature tolerances, which may affect community composition and ecosystem functions more profoundly than a gradual long-term temperature increase (Vasseur *et al*. 2014; Stillman, 2019). Heatwaves have been shown to cause biodiversity loss in both marine and freshwater habitats (Brauko *et al*. 2020; Sabater *et al*. 2022). Particularly for eukaryotic microorganisms, heat stress has resulted in significant shifts in community composition (Hao *et al*. 2018; Thomson *et al*. 2019). However, it is assumed that short-term disturbances like heatwaves are succeeded by complete or partial recovery (Bender *et al*. 1984; Harris *et al*. 2018).

Colorless heterotrophic nanoflagellates play an important role in regulating the quantity and biomass of bacterial populations and profoundly influence the morphological and taxonomic structure of bacterial communities (Sherr & Sherr 2002). Consequently, alterations in both the quantity and quality of bacterivory, resulting from shifts in the flagellate community, can influence the overall structure of aquatic food webs. The Chyrosphytes are among the most important bacterivorous grazers in freshwater habitats (Finlay & Esteban 1998). They are widely distributed across various habitats, primarily inhabiting freshwaters but also extending into terrestrial and marine ecosystems (Kristiansen and Škaloud 2017). They further encompass a broad range of different morphologies (Škaloud *et al*. 2013; Škaloudová & Škaloud 2013). Interestingly, molecular analysis demonstrated high molecular diversity behind distinct morphological forms (Grossmann *et al*. 2016). This cryptic diversity likely resulted from parallel evolution, wherein similar morphological forms evolved independently in different lineages (Graupner *et al*. 2018). The ecological significance of this phenomenon, however, remains unclear. The morphology of flagellates has been shown to correlate with various aspects of predator-prey interactions, including preferences for food size and feeding mechanisms (Boenigk & Arndt 2000 a, b). Consequently, flagellates exhibiting similar morphologies can feed on the same prey, thereby serving equivalent roles in the food web. On the other hand, there is no correlation between morphology and adaptations to environmental abiotic factors. In the order Ochromonadales, significant differences in temperature tolerance were observed among clades with similar morphologies (Boenigk *et al*. 2007; Nolte *et al*. 2010). Such clade-specific responses to changing environmental conditions can result in a shift of the community composition (Boden *et al*. 2023). This suggest that cryptic diversity might disguise a turn-over of cryptic taxa that are differently adapted to abiotic factors, thereby stabilizing the structure of food webs and ecosystem functions during the occurrence of stressors despite significant changes in the taxon composition (Boden *et al*. 2023).

Here, we investigate the effects of heatwaves on the growth of three cryptic taxa associated with *Pedospumella, Spumella*, and *Poteriospumella*. Given the variation in temperature tolerance, we anticipate that the three strains will respond differently to the temperature increase. Additionally, we expect the strains to grow differently in mixed cultures compared to the single-strain cultures because biotic interactions between the strains will influence the growth of each taxon. We predict more pronounced fluctuations in the total abundance of flagellates in single-strain cultures compared to mixed cultures because the presence of multiple taxa with varying temperature tolerances in mixed cultures may buffer the effects of the heatwave. Consequently, we also expect less fluctuation in the total abundance of prey bacteria in mixed cultures, as the presence of multiple flagellate strains feeding on the same prey should stabilize the predator-prey interaction and ensure functional redundancy in the face of environmental changes and potential shifts in the flagellate community.

## Materials and Methods

### Strains & Cultivation

The strains *Pedospumella encystans* JBM/S11 (Boenigk & Findenig, 2010), *Spumella rivalis* AR4A6 (Boenigk & Findenig, 2010), and *Poteriospumella lacustris* JBM10 (Boenigk & Findenig, 2010) were previously isolated from soil or freshwater samples originating from different geographical locations (Table 1). JBM/S11 and AR4A6 are routinely grown as xenic cultures in inorganic basal medium (Hahn *et al*. 2003), using wheat grains as a food source, in cell culture flasks (25 cm^3^ with filter screw cap, TTP Techno Plastic Products AG), while JBM10 is cultivated as an axenic culture in NSY medium (Nutrient broth, Peptone from soybean, Yeast extract; Hahn *et al*. 2003) in 100 mL Erlenmeyer flasks. All strains are grown in a climate chamber (SANYO Electric Co. Ltd., Osaka, Japan) at 15 °C under a 14h:10h light-dark cycle. The bacterial strain *Linmohabitans* spp. IID5 is grown in NSY medium in 100 mL Erlenmeyer flasks at room temperature (RT) with constant shaking (96 rpm, orbital shaker, LAUDA-Brinkmann, LP, New Jersey, USA).

**Table 1.** Origin and affiliation of strains used in this study.

| Strain | Species designation | Geographical Origin | Habitat origin | Media |
| --- | --- | --- | --- | --- |
| JBM/S11 | *Pedospumella encystans* | Austria, Mondsee, near Rauchhaus | soil | IB + grain of wheat |
| AR4A6 | *Spumella rivalis* | Austria, River Fuschler Ache | freshwater | IB + grain of wheat |
| JBM10 | *Poteriospumella lacustris* | Austria, Mondsee | freshwater | NSY (3g/L) |

### Experimental set-up with single-strain and mixed cultures

Flagellates were harvested by centrifugation for 10 min at 2820 g at RT to remove the original medium and each resulting pellet was then resuspended in 50 mL IB medium. One mL aliquots of a *Linmohabitans* spp. IID5 culture were centrifuged for 10 minutes at 15000 g at RT and, after removal of the supernatant, resuspended in IB medium. One milliliter of the washed bacteria was added to each flagellate culture as food source. These pre-cultures were then incubated at 23 °C with a 14h:10h light-dark cycle for three days to acclimatize to the new conditions. The bacteria from both xenic strains were isolated through filtration (diameter 25 mm, pore size 0.45 μm, Millipore GTTP 02500, Eschborn, Germany) and cultivated in NSY medium in a 100 mL Erlenmeyer flask at 15 °C with a 14h:10h light-dark cycle. All experiments were conducted in triplicates in 75 mL of IB medium in 100 mL Erlenmeyer flasks. The flagellate abundance was adjusted to approximately 30000 cells per mL in single-strain cultures. In the mixed cultures, the three strains were mixed to achieve relative abundances of 20 % *Pedospumella*, 30 % *Poteriospumella*, and 50 % *Spumella*, mimicking the relative abundances of the respective clades in natural environments during late spring (Nolte *et al*. 2010). The concentration of food bacteria was set to 2 × 10^7^ bacteria per mL, consisting predominately of *Limnohabitans* spp. IID5 in addition to 10000 cells per mL of the previously cultivated bacteria from the xenic flagellate cultures to ensure uniformed bacterial backgrounds in all cultures. Further, wheat grains were preheated at 70° C for one hour and one wheat grain was added to each culture as food source for the bacteria. All cultures were initially incubated at 23 °C with a 14h:10h light-dark cycle for two days. The temperature was then increased to 27 °C for three days before being reduced back to 23 °C for a two-day recovery phase. Subsamples were collected daily and fixed with 2% paraformaldehyde (PFA) for 1 hour at RT or overnight at 4 °C, and 4 mL of each fixed sample were filtered onto white polycarbonate filters (diameter 25 mm, pore size 0.2 μm, Millipore GTTP 02500, Eschborn, Germany). The filters were air-dried and stored at -20 °C for further use.

### Determination of cell counts and growth rates in single-strain and mixed cultures

Sections of the filters were cut out with a razor blade. To determine total flagellate abundance in single-strain cultures, the filter sections were stained with 4,6-diamidino-2-phenylindole (DAPI; 0.1 mg/ml) for 10 minutes at RT. To remove non-specific staining, the filters were washed in sterile distilled water for 1 minute at RT, then dehydrated with 100% ethanol for several seconds at RT and air-dried. To determine the fraction of each strain in the mixed cultures, *Pedospumella encystans, Spumella rivalis*, and *Poteriospumella lacustris* were stained using fluorescently labelled clade-specific probes (O1C531, O2C613, O3C723; Boden *et. al* 2023; 50 ng DNA/μL, Eurofins Genomics Germany GmbH, Ebersberg, Germany). Therefore, filter sections were incubated sequentially in 50%, 80%, and 100% ethanol solutions to dehydrate the samples. For each dehydration step, the filters were incubated for 3 minutes at RT. Then, the cells were hybridized with Cy3-labeled single-strand oligonucleotide probes as described by Glöckner *et al*. (1996). To optimize signal detection rates, the concentration of SDS was increased to 0.02 % in the hybridization buffer, and the hybridization time was extended to 3 hours at 46 °C. To also determine total flagellate abundance in the mixed cultures, the filter sections were afterward counterstained with DAPI as described before. After staining, the filter sections were mounted in the non-hardening and non-bleaching mounting medium CitiFluorTM AF2 (Citifluor, Ltd., London, United Kingdom), and images were captured using the NIS-Elements BR imaging software (Nikon Corp., Tokyo, Japan) and a Nikon Eclipse 80i microscope (Nikon Corp., Tokyo, Japan) with a 100x objective and appropriate fluorescence filters to detect DAPI and Cy3 signals. DAPI and Cy3 signals were counted manually in Fiji (Schindelin *et al*. 2012).

Growth rates were calculated for three distinct phases throughout the experiment using R (v4.3.3, R Core Team 2024) and R Studio (v2023.12.1, RStudio Team, 2024). Specifically, growth rates were determined during the initial acclimatization phase from exponentially growing single strains as well as for all flagellates in the mixed cultures. Further, growth rates were assessed at the end of the heatwave, i.e., between days five and six, and during the recovery phase, i.e., between days seven and eight. Cell abundances, growth rates and the relative share of each taxa in the mixed cultures were plotted using R, R Studio and the R packages “readxl”, “gglot”, “cowplot” and “dplyr” (Wickham and Bryan 2023; Wickham 2016; Wilke 2024; Wickham *et al*. 2023)

### Statistical data analysis

Statistical data analysis was performed in R and R Studio. Growth rates were statistically compared among each other via Welch’s ANOVA and Games-Howell post-hoc test using the R-packages “readxl”, “dplyr” and “rstatix” (Wickham and Bryan 2023; Wickham *et al*. 2023; Kassambara 2023). Similarly, the relative shares of *Pedospumella encystans, Spumella rivalis* and *Poteriospumella lacustris* in the mixed cultures at different sampling days were statistically compared via Welch’s ANOVA and Games-Howell post-hoc test. Variability in bacterial abundances was statistically compared among all cultures via Brown-Forsythe test.

## Results

After a lag phase between day one and day two, an increase in total flagellate abundance starting from day two was observed in all cultures (Figure 1). Growth occurred between day two and day four in all cultures, with the exception of *Poteriospumella* single-strain experiments, where growth ceased by day three (Table 1).

**Figure 1.**
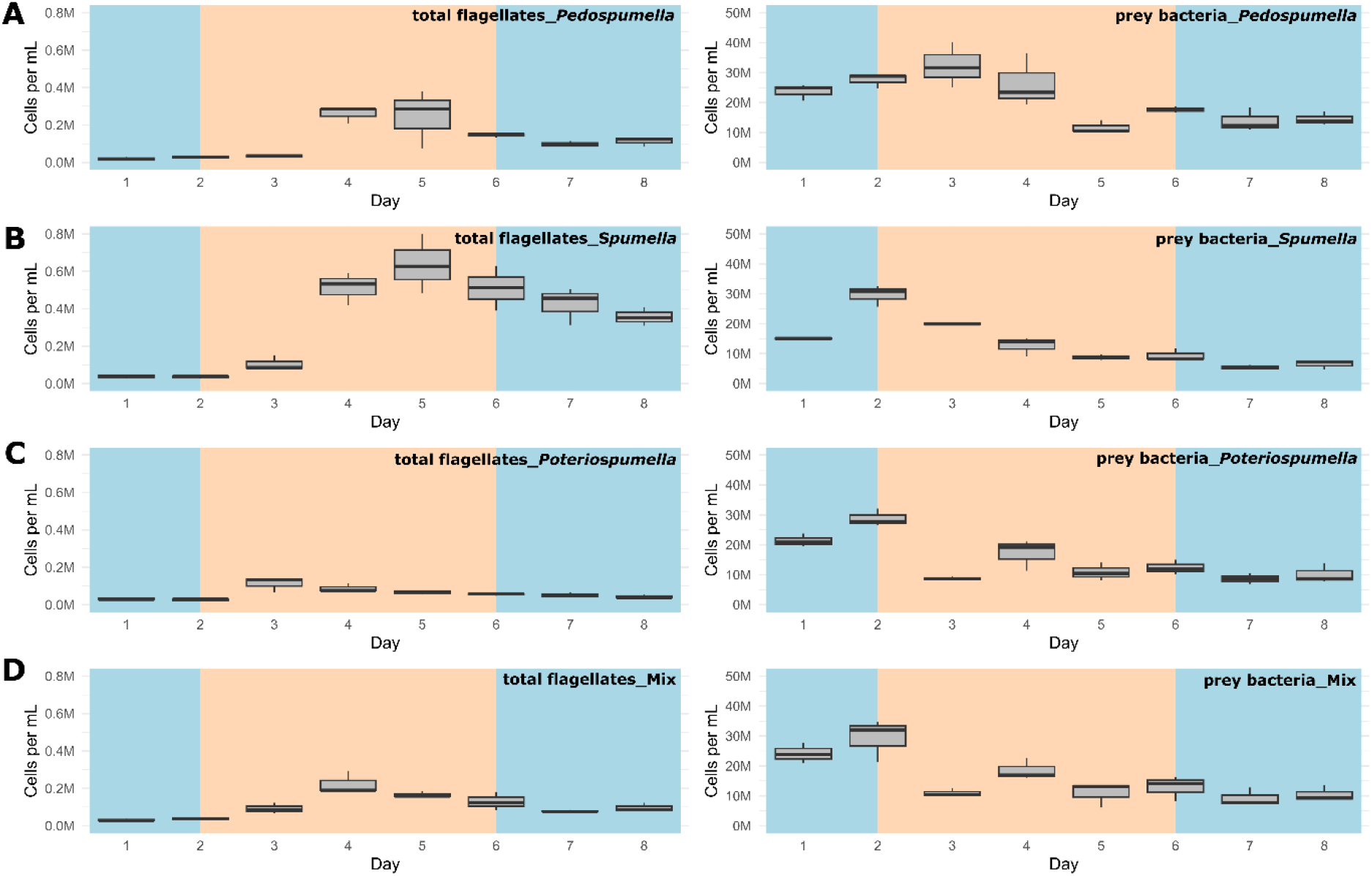
Total absolute abundance of flagellates (left) and bacteria (right) during phases of normal (blue) and increased (orange) temperature in single-stain cultures of A) *Pedospumella encystans* (JBMS11), B) *Spumella rivalis* (AR4A6), C) *Poteriospumella lacustris* (JBM10) and D) mixed cultures containing all three strains.

During this initial growth phase, hereafter referred to as “initial phase”, significant differences among the cultures were observed when comparing flagellate growth in the single-strain cultures and in the mixed cultures (p = 0.04). Growth rates of the taxa were not significantly different when grown alone (p = 1.00; Figure 2). But growth rates significantly differed during the initial phase (p = 0.004) between flagellates growing in single-strain culture and the same taxon growing in mixed culture: Growth rates for *Spumella* (p = 0.02) and *Pedospumella* (p = 0.02) were significantly lower in the mixed cultures compared to when they were grown individually. In contrast, growth rates of *Poteriospumella* did not differ between treatments (p = 1.00; Figure 2). During the heatwave total flagellate abundance decreased in all cultures (Figure 1) and no significant differences were observed between the three taxa (p = 0.98). Similarly, no significant differences were observed when comparing growth of the three taxa in single-strain culture and in mixed culture (*Pedospumella*: p = 1.00; *Spumella*: p = 1.00; *Poteriospumella*: p = 1.00; Figure 2). During the recovery phase no significant differences were observed in total flagellate growth when comparing single-strain experiments and mixed cultures (p = 0.37). Interestingly, the growth rates of *Poteriospumella* and *Spumella* in single-strain cultures were slightly but not significantly lower than in the mixed cultures (*Poteriospumella*: p = 0.68; *Spumella*: p = 0.64).

**Figure 2.**
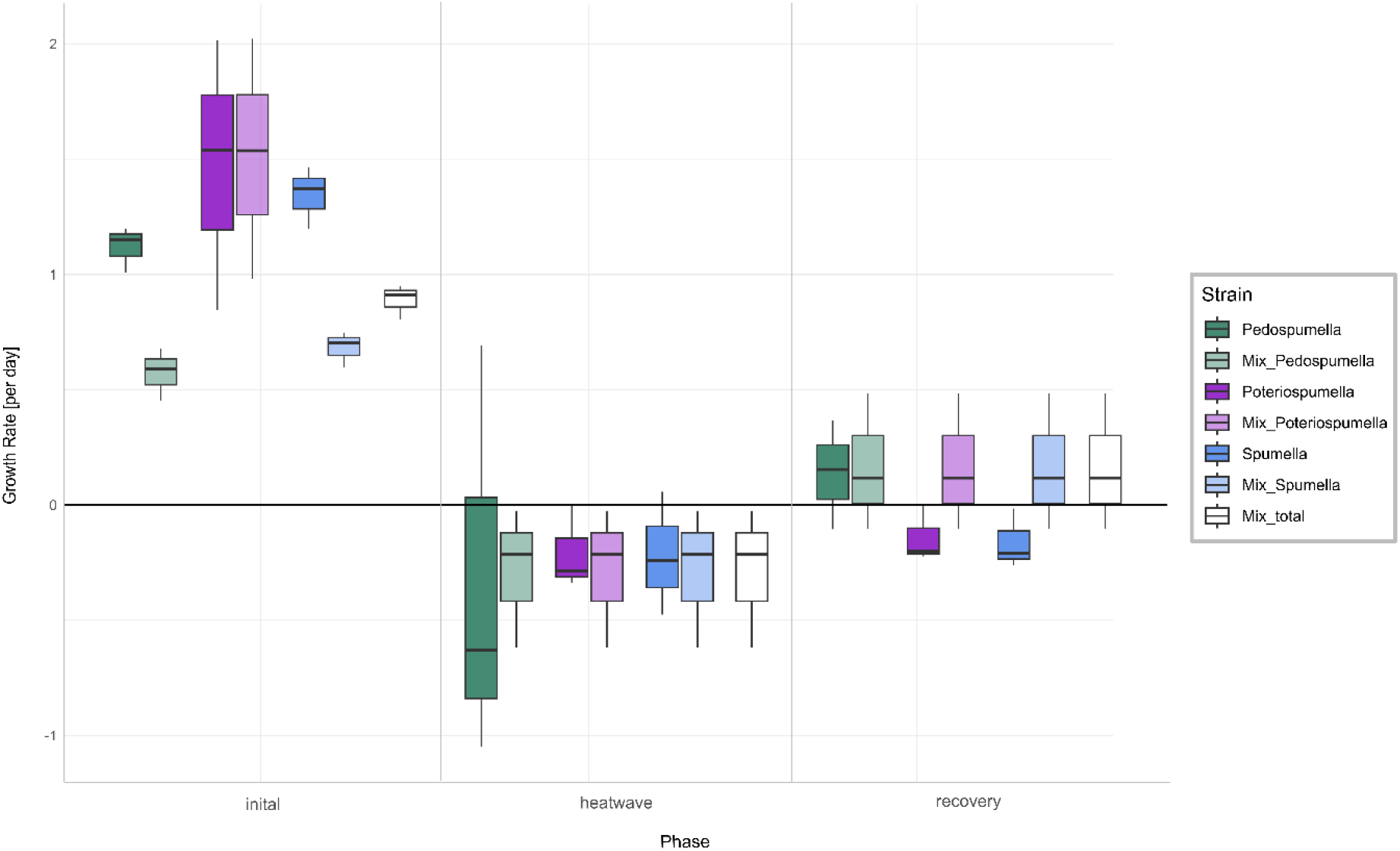
Growth rates [day–1] of *Pedospumella encystans, Spumella rivalis* and *Poteriospumella lacustris* in single-strain cultures and in mixed cultures as well as total flagellate growth in the mixed cultures during the initial phase of the experiment, at the end of the heatwave and during the recovery phase. Negative growth rates correspond to mortality rates.

The relative share of all three taxa in the mixed culture (Figure 2) changed significantly during the onset of the heatwave but was stable before and after: No significant changes were observed for any of the three taxa during the initial acclimatization phase, i.e., between day one and day two (*Pedospumella*: p = 1.00; *Spumella*: p = 0.20; *Poteriospumella*: p = 0.48). During the onset of the heatwave, i.e., between day two and three, the relative share of *Spumella* decreased significantly from 52 % on day two to 34 % on day three (p < 0.001). Similarly, the relative share of *Pedospumella* decreased significantly from 21 % on day two to 11 % on day three (p < 0.001). In contrast, the share of *Poteriospumella* increased significantly from 29 % on day two to 55 %, dominating the community on day three (p < 0.001). After this shift in community composition during onset of the heatwave, no further significant changes of the relative share were observed both during the heatwave (*Pedospumella*: p = 0.97; *Spumella*: p = 0.99; *Poteriospumella*: p = 0.96) and during the following recovery phase (*Pedospumella*: p = 0.95; *Spumella*: p = 1.00; *Poteriospumella*: p = 1.00). On day seven, *Poteriospumella* still dominated the community, constituting on average 56 % of the total flagellate abundance, while *Pedospumella* and *Spumella* represented 12 % and 34 %, respectively, of all flagellates in the mixed cultures.

Furthermore, variability of the bacterial abundances over time differed significantly among the cultures (p = 0.01). The relative variability in bacterial abundances was assessed by calculating the coefficient of variation (CV) for each culture. *Pedospumella* single-strain cultures exhibited the lowest CV at 39.1 %, indicating the least variability in bacterial abundance throughout the experiment. In contrast, *Spumella* single-strain cultures showed the highest CV at 58.8 %. *Poteriospumella* single-strain cultures and mixed cultures (mL) displayed similar CVs of 49.4 % and 49.6 %, respectively.

## Discussion

The impact of temperature shifts, e.g. due to global warming and heatwaves, has been the subject of numerous studies (Font *et al*. 2021; Woolway *et al*. 2021), revealing substantial shifts in protist communities (Hao *et al*. 2018; Thomson *et al*. 2019). We observed significant differences both among the different cultures and when comparing growth of each taxa in single-strain and mixed cultures (Figure 2). While total flagellate growth was generally higher in single-strain experiments than in the mixed cultures, *Poteriospumella* exhibited the highest growth rate among all cultures. Interestingly, similar growth rates were observed for *Poteriospumella* in both single-strain and mixed cultures, contrasting with *Pedospumella* and *Spumella*, which displayed significantly lower growth rates in mixed cultures compared to when they were grown individually (Figure 2). This is in accordance with previous research that identified *Poteriospumella*’s competitive advantage, particularly under elevated temperature conditions (Boden *et al*. 2023). Additionally, we observed a shift in the community composition in the mixed cultures. At the beginning of the experiment, the community was dominated by *Spumella* but its relative share decreased significantly during the onset of the heatwave. In contrast, the relative abundance of *Poteriospumella* increased significantly, becoming the dominant taxa approximately 24 hours after the temperature rise (Figure 3). Similar temperature-induced shifts in relative abundance have been observed in seasonal environmental data, with *Spumella* typically dominating in winter and *Poteriospumella* in summer (Nolte *et al*. 2010).

**Figure 3.**
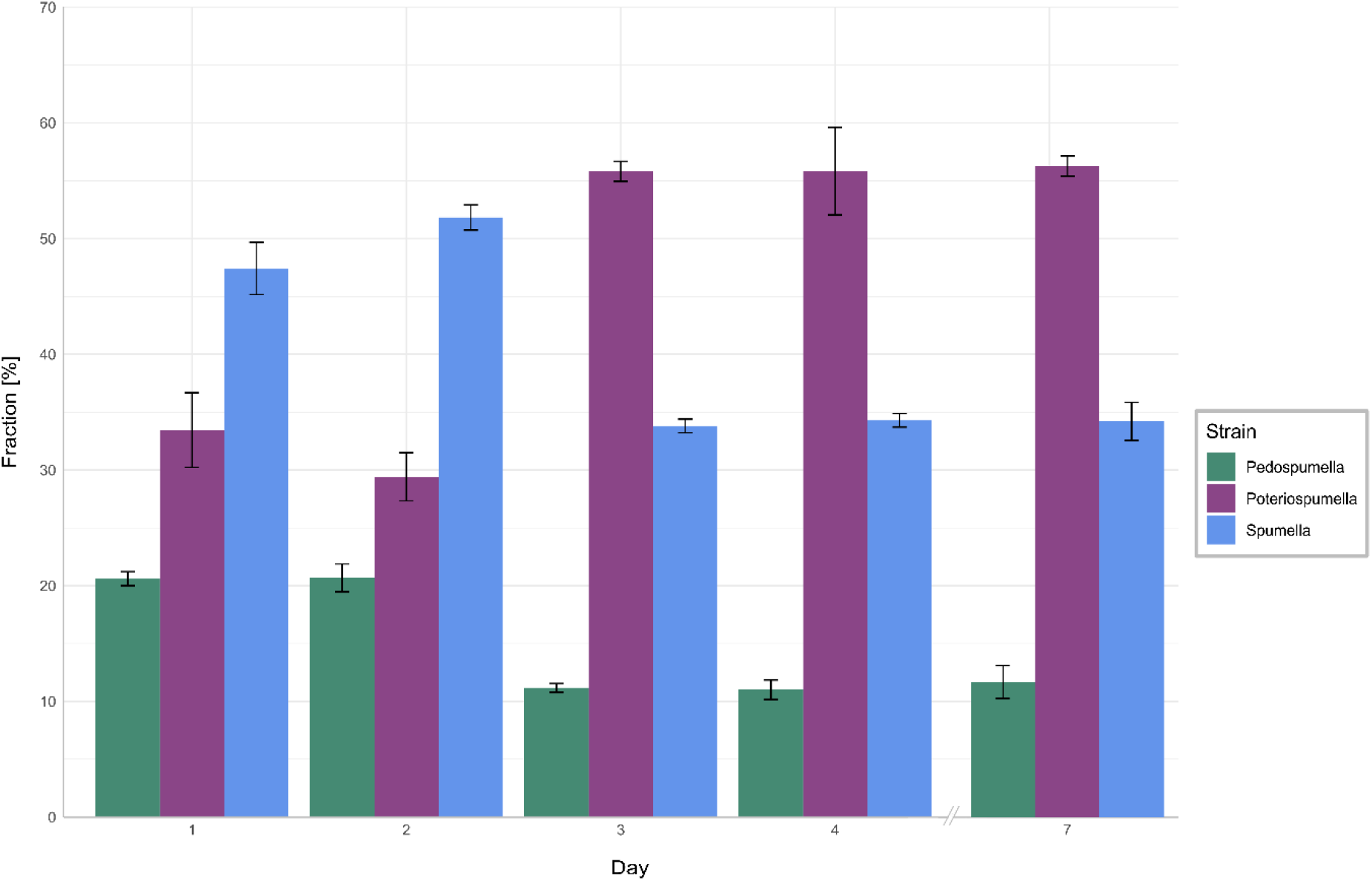
Relative shares of *Pedospumella encystans, Poteriospumella lacustris* and *Spumella rivalis* in the mixed cultures. The bars indicate the mean percentage of each strain in all mixed cultures on day 1, day 2, day 3, day 4, and day 7. Error bars represent standard deviations.

No shifts in community composition were observed at the end of the heatwave and during the recovery phase, i.e., the community composition remained unchanged despite changing temperature conditions (Figure 3). This was unexpected as the influence of abiotic factors such as temperature, salinity or oxygen availably on flagellate abundance and community composition has been demonstrated in previous studies (Nolte *et al*. 2010; Princiotta and Sanders, 2017; Gran-Stadniczeñko *et al*. 2018). However, most of these studies were investigating the effects of abiotic factor in environmental samples, indicating that mock communities in laboratory experiments may be less dynamic than natural communities, possibly due to reduced competition in species-poor communities. At the end of the heatwave, the growth rates of *Poteriospumella*, both in single-strain and mixed cultures, were similar to those of *Pedospumella* and *Spumella* and no competitive advantage was identified for any of the three taxa during that later phase. The reasons why *Poteriospumella* failed to maintain its competitive advantage throughout the entire duration of the heatwave remain unclear. All cultures reached maximal abundance latest by day five, therefore potential reasons include density-dependent factors like the accumulation of toxic metabolic byproducts such as acids and alcohols, or oxygen depletion, which are common phenomena in cultures with high cell density and would affect all three taxa equally (Rouf *et al*. 2017). Beside abiotic factors, biotic interactions can also influence flagellate growth in the cultures. High cell densities can also lead to cannibalism among flagellates, particularly when alternative prey is scarce or competition is high (Martel and Flynn 2008; Yang *et al*. 2020). However, we found no indications of cannibalism in our samples, likely because the concentration of prey bacteria remained above 5 x 10^6^ cells / mL throughout the experiment, which was sufficiently high to prevent starvation (Boenigk *et al*. 2002). Given the consistently high concentration of prey bacteria, it is unlikely that nutrient depletion, a common cause of mortality in dense microbial cultures (Rouf *et al*. 2017), was responsible for the decline in total flagellate abundances at the end of the heatwave. Our results indicate that abiotic and/or biotic density-dependent factors overshadow the effects of the temperature increase during the later half of the heatwave. While heterotropic nanoflagellates can reach abundances of several thousand cells per mL in natural habitats, their total flagellate abundances remain notably lower than the maximum abundances that were reached in this study (Boenigk and Arndt, 2002). Therefore, the response to the temperature increase that was observed in the initial phase may be a closer representation of natural habitats than towards the end to the heatwave. In the recovery phase, growth rates of *Spumella* and *Poteriospumella* were slightly but not significantly higher in the mixed cultures than in single-strain cultures, indicating that the lower total flagellate abundance during the recovery phase may reduce the impact of density-dependent factors, thereby allowing temperature-related effects to become dominant again.

As important players in the top-down control of bacterial populations, changes in both the total abundance and community composition of heterotrophic nanoflagellates significantly influence the dynamics of bacterial prey populations (Sherr and Sherr, 2002). In this study, variability in the bacterial abundances differed among the single strain-cultures, with the highest variation observed in *Spumella* single-strain. Notably, within mixed cultures, bacterial abundances displayed similar variability than in the *Poteriospumella* single-strain cultures. The strong fluctuations observed in *Spumella* single-strain cultures were likely mitigated in the mixed cultures by the presence and dominance of *Poteriospumella*. These results further support the hypothesis that functional redundancy, demonstrated here by the simultaneous presence of taxa capable of feeding on the same prey but exhibiting different adaptations to abiotic factors, can stabilize habitats when confronted with changing environmental conditions. In conclusion, our findings highlight the complexities of protist community dynamics under thermal stress, indicating that while temperature can drive significant changes in abundance and composition of flagellate communities, other factors such as density-dependent interactions and prey availability also significantly impact microbial communities. Our study indicates that stabilizing effects due to functional redundancy may be pronounced as long as density-dependent factors play a minor role. This study underscores the importance of considering both abiotic and biotic factors when assessing the impacts of climate change on microbial communities, as well as the potential for functional redundancy to stabilize ecosystems facing environmental fluctuations.

## Acknowledgements

This study was performed within the Collaborative Research Center (CRC) RESIST and analyses were mainly done in Project A06, funded by the German Research Foundation (DFG) – CRC 1439/1; project number INST 20876/402–1. We acknowledge support by the Open Access Publication Fund of the University of Duisburg–Essen

## References

Bender, E.A., Case, T.J. and Gilpin, M.E. (1984), Perturbation Experiments in Community Ecology: Theory and Practice. Ecology, 65: 1–13. 10.2307/1939452

Bernhardt JR, O’Connor MI,Sunday JM, Gonzalez A. 2020 Life influctuating environments. Phil. Trans. R. Soc. B375: 20190454. 10.1098/rstb.2019.0454

Boden, L., Sieber, G. and Boenigk, J. (2023): Effects of stressors on growth and competition between different cryptic taxa affiliated with Ochromonadales (Chrysophyceae). Fottea. 23. 235–245. 10.5507/fot.2023.003.

Boenigk, J. & Arndt, H. (2000): Particle handling during interception feeding by four species of heterotrophic nanoflagellates. – J. Euk. Mic. 47: 350–358.

Boenigk J, Arndt H. Bacterivory by heterotrophic flagellates: community structure and feeding strategies. Antonie Van Leeuwenhoek. 2002 Aug;81(1-4):465–80. doi: 10.1023/a:1020509305868. PMID: 12448743.

Boenigk, J. & Arndt, H. (2000): Comparative studies on the feeding behavior of two heterotrophic nanoflagellates: the filter–feeding choanoflagellate Monosiga ovata and the raptorial–feeding kinetoplastid Rhynchomonas nasuta. – Aquat. Microb. Ecol. 22: 243–249.

Boenigk, J., Matz, C., Jürgens, K., & Arndt, H. (2002). Food concentration-dependent regulation of food selectivity of interception-feeding bacterivorous nanoflagellates. Aquatic Microbial Ecology, 27(2), 195–202.

Boenigk, J.; Jost, S.; Stoeck, T. & Garstecki, T. (2007): Differential thermal adaptation of clonal strains of aprotist morphospecies originating from different climatic zones. – Environ. Microbiol. 9: 593–602.

K. M. Brauko, A. Cabral, N. V. Costa, J. Hayden, C. E. P. Dias, E. S. Leite, R. D. Westphal, C. M. Mueller, J. M. Hall-Spencer, R. R. Rodrigues, L. R. Rörig P. R. Pagliosa, A. L. Fonseca, O. E. Alarcon, and P. A. Horta, “Marine heatwaves, sewage and eutrophication combine to trigger deoxygenation and biodiversity loss: A sw atlantic case study,” Frontiers in Marine Science, vol. 7, 12 202

Biggs, C., Yeager, L., Bolser, D., Bonsell, C., Dichiera, A., Hou, Z., Keyser, S., Khursigara, A., Lu, K., Muth, A., Negrete, B., & Erisman, B. (2020). Does functional redundancy affect ecological stability and resilience? A review and meta-analysis. Ecosphere. 10.1002/ecs2.3184.

Chen, H., Ma, K., Lu, C., Fu, Q., Qiu, Y., Zhao, J., Huang, Y., Yang, Y., Schadt, C., & Chen, H. (2022). Functional Redundancy in Soil Microbial Community Based on Metagenomics Across the Globe. Frontiers in Microbiology, 13. 10.3389/fmicb.2022.878978.

Díaz-Almeyda Erika M., Prada C., Ohdera A. H., Moran H., Civitello D. J., Iglesias-Prieto R., Carlo T. A., LaJeunesse T. C. and Medina M. 2017. Intraspecific and interspecific variation in thermotolerance and photoacclimation in Symbiodinium dinoflagellates. Proc. R. Soc. B. 28420171767. 10.1098/rspb.2017.1767

Finlay, B.J. & Esteban, G.F. (1998): Freshwater protozoa: biodiversity and ecological function. – Biodivers. Conservation 7: 1163–1186.

Font, R., Khamis, K., Milner, A., Smith, G., & Ledger, M. (2021). Low flow and heatwaves alter ecosystem functioning in a stream mesocosm experiment. The Science of the total environment, 777, 146067. 10.1016/j.scitotenv.2021.146067.

Gran-Stadniczeñko, S., Egge, E., Hostyeva, V., Logares, R., Eikrem, W., & Edvardsen, B. (2018). Protist Diversity and Seasonal Dynamics in Skagerrak Plankton Communities as Revealed by Metabarcoding and Microscopy. The Journal of Eukaryotic Microbiology, 66, 494–513. 10.1111/jeu.12700.

Graupner, N.; Jensen, M.; Bock, C.; Marks, S.; Rahmann, S.; Beisser, D. & Boenigk, J. (2018): Evolution of heterotrophy in chrysophytes as reflected by comparative transcriptomics. – FEMS Microbiol. Ecol 94: fiy039. DOI: 10.1093/femsec/fiy039

Grossmann, L., Bock, C., Schweikert, M., & Boenigk, J. (2016). Small but Manifold – Hidden Diversity in “Spumella-like Flagellates”. The Journal of Eukaryotic Microbiology, 63, 419–439. 10.1111/jeu.12287.

Glöckner, F.O.; Amann, R.; Alfreider, A.; Pernthaler, J.; Psenner, R.; Trebesius, K. & Schleifer, K.–H. (1996): An in situ hybridization protocol for detection and identification of planktonic bacteria. – Syst. Appl. Microbiol. 19: 403–406.

Hahn, M.W.; Lundsdorf, H.; Wu, Q.L.; Schauer, M.; Hofle, M.G.; Boenigk, J. & Stadler, P. (2003): Isolation of novel ultramicrobacteria classified as actinobacteria from five freshwater habitats in Europe and Asia. – Appl. Environ. Microb. 69: 1442–1451

Hao, B.; Roejkjaer, A.F.; Wu, H.; Jeppesen, E.; Li, W. (2018): Responses of primary producers in shallow lakes to elevated temperature: a mesocosm experiment during the growing season of Potamogeton crispus. – Aquat. Sci. 80: 34

Harris, R.M.B., Beaumont, L.J., Vance, T.R. et al. Biological responses to the press and pulse of climate trends and extreme events. Nature Clim Change 8, 579–587 (2018).

IPBES (2019): Global assessment report on biodiversity and ecosystem services of the Intergovernmental Science-Policy Platform on Biodiversity and Ecosystem Services. E. S. Brondizio, J. Settele, S. Díaz, and H. T. Ngo (editors). IPBES secretariat, Bonn, Germany. 1148 pages. 10.5281/zenodo.3831673

IPCC. 2021: Summary for Policymakers. In: Climate Change 2021: The Physical Science Basis. Contribution of Working Group I to the Sixth Assessment Report of the Intergovernmental Panel on Climate Change [Masson-Delmotte, V., P. Zhai, A. Pirani, S. L. Connors, C. Péan, S. Berger, N. Caud, Y. Chen, L. Goldfarb, M. I. Gomis, M. Huang, K. Leitzell, E. Lonnoy, J.B.R. Matthews, T. K. Maycock, T. Waterfield, O. Yelekçi, R. Yu and B. Zhou (eds.)]. Cambridge University Press. In Press. Kassambara A (2023). _rstatix: Pipe-Friendly Framework for Basic Statistical Tests_. R package version 0.7.2, <https://CRAN.R-project.org/package=rstatix>.

Kristiansen, J. & Škaloud, P. (2017): Chrysophyta. In: Archibald, J.M.; Simpson, A.G.B. & Slamovits, C.H. (eds.): Handbook of the Protists. – pp. 331–366, Springer

Lau, G. X. H., Fan, H. Y., Halim, M. A., Cheah, Y. K., Wong, C. M. V. L. 2024. Forecasting the impact of global warming on soil bacterial communities using simulated systems. Malaysian Journal of Microbiology, Vol 20(3) pp. 397–405. DOI: 10.21161/mjm.230323

Maire, J., Buerger, P., Chan, W., Deore, P., Dungan, A., Nitschke, M., & Oppen, M. (2022). Effects of Ocean Warming on the Underexplored Members of the Coral Microbiome. Integrative and Comparative Biology, 62, 1700–1709. 10.1093/icb/icac005.

H. Mooney, A. Larigauderie, M. Cesario, T. Elmquist, O. Hoegh-Guldberg, S. Lavorel, G.M. Mace, M. Palmer, R. Scholes, T. Yahara Biodiversity, climate change, and ecosystem services Curr. Opin. Environ. Sustain., 1 (Issue 1) (2009), pp. 46–54, 10.1016/j.cosust.2009.07.006

Jain, R., & Saraf, M. (2021). EXPLORING THE ABIOTIC AND BIOTIC STRESS TOLERANCE POTENTIAL OF RHIZOBACTERA ISOLATED FROM CYAMOPSIS. Journal of Advanced Scientific Research. 10.55218/jasr.202112327.

Martel, C., & Flynn, K. (2008). Morphological controls on cannibalism in a planktonic marine phagotroph.. Protist, 159 1, 41–51. 10.1016/J.PROTIS.2007.05.003.

Mascarin, G., Pereira-Junior, R., Fernandes, É., Quintela, E., Dunlap, C., & Arthurs, S. (2018). Phenotype responses to abiotic stresses, asexual reproduction and virulence among isolates of the entomopathogenic fungus Cordyceps javanica (Hypocreales: Cordycipitaceae). Microbiological research, 216, 12–22. 10.1016/j.micres.2018.08.002.

Naeem, S. 1998. Species redundancy and ecosystem reliability. Conservation Biology 12:39–45.

Nolte, V.; Pandey, R.V.; Jost, S.; Medinger, R.; Ottenwaelder, B.; Boenigk, J. & Schlötterer, C. (2010): Contrasting seasonal niche separation between rare and abundant taxa conceals the extend of protist diversity. – Mol. Ecol. 19: 2908–2915.

Princiotta, S., & Sanders, R. (2017). Heterotrophic and mixotrophic nanoflagellates in a mesotrophic lake: Abundance and grazing impacts across season and depth. Limnology and Oceanography, 62. 10.1002/lno.10450.

Proesmans, W., Andrews, C., Gray, A., Griffiths, R., Keith, A., Nielsen, U. N., Spurgeon, D., Pywell, R., Emmett, B., & Vanbergen, A. J. (2022). Long-term cattle grazing shifts the ecological state of forest soils. Ecology and Evolution, 12, e8786. 10.1002/ece3.8786

R Core Team (2024). R: A Language and Environment for Statistical Computing. R Foundation for Statistical Computing, Vienna, Austria. <https://www.R-project.org/>.

A Rouf, Varsha Kanojia, HR Naik, Bazilla Naseer and Tahiya Qadri. An overview of microbial cell culture. J Pharmacogn Phytochem 2017;6(6):1923–1928.

RStudio Team (2024). RStudio: Integrated Development Environment for R. RStudio, PBC, Boston, MA. URL http://www.rstudio.com/.

Ruokolainen L, Lindén A, Kaitala V, Fowler MS. 2009 Ecological and evolutionary dynamics under coloured environmental variation. Trends Ecol. Evol. 24, 555–563. (doi: 10.1016/j.tree.2009.04.009)

S. Sabater, A. Freixa, L. Jiméenez J. López-Doval, G. Pace, C. Pascoal, N. Perujo, D. Craven, and J. D. González-Trujillo, “Extreme weather events threaten biodiversity and functions of river ecosystems: evidence from a meta-analysis,” Biological Reviews, 10 20

Schulhof MA, Allen AE, Allen EE, et al. Sierra Nevada mountain lake microbial communities are structured by temperature, resources and geographic location. Mol Ecol. 2020; 29: 2080–2093. 10.1111/mec.15469

Schindelin, J., Arganda-Carreras, I., Frise, E., Kaynig, V., Longair, M., Pietzsch, T., … Cardona, A. (2012). Fiji: an open-source platform for biological-image analysis. Nature Methods, 9(7), 676–682. doi:10.1038/nmeth.2019

Škaloud, P.; kristiansen, J. & Škaloudová, m. (2013): Developments in the taxonomy of silica–scaled chrysophytes – from morphological and ultrastructural to molecular approaches. – Nord. J. Bot. 31: 385–402.

Škaloudová, m. & Škaloud, P. (2013): A new species of Chrysosphaerella (Chrysophyceae: Chromulinales), Chrysosphaerella rotundata sp. nov., from Finland. – Phytotaxa 130: 34–42

Sherr, E.B. & Sherr, B.F. (2002): Significant of predation by protists in aquatic microbial food webs. – Antonie Van Leewenhoek 81: 293–308.

Stillman, J. H. (2019). Heat waves, the new normal: Summertime temperature extremes will impact animals, ecosystems, and human communities. Physiology, 34(2), 86–100. 10.1152/physiol.00040.2018

Sun, X. & Arnott, S.E. (2022): Interactive effects of increased salinity and heatwaves on freshwater zooplankton communities in simultaneous and sequential treatments. – Freshw. Biol. 67: 1604–1617

Thomson, A.H. & Manoylov, K.M. (2019): Algal community dynamics within the Savannah River estuary, Georgia under anthropogenic stress. – Estuar. Coasts 42: 1459–1474

Urrutia-Cordero, P., Ekvall, M. K., Ratcovich, J., Soares, M., Wilken, S., Zhang, H., & Hansson, L. A. (2017). Phytoplankton diversity loss along a gradient of future warming and brownification in freshwater mesocosms. Freshwater Biology, 62(11), 1869–1878. 10.1111/fwb.13027

Vasseur, D. A., DeLong, J. P., Gilbert, B., Greig, H. S., Harley, C. D. G., McCann, K. S., Savage, V., Tunney, T. D., & O’Connor, M. I. (2014). Increased temperature variation poses a greater risk to species than climate warming. Proceedings of the Royal Society B: Biological Sciences, 281(1779), 20132612. 10.1098/rspb.2013.2612

Walker, B. H. 1992. Biodiversity and ecological redundancy. Conservation Biology 6:18–23.

H. Wickham. ggplot2: Elegant Graphics for Data Analysis. Springer-Verlag New York, 2016.

Wickham H, Bryan J (2023). _readxl: Read Excel Files_. R package version 1.4.3, <https://CRAN.R-project.org/package=readxl>.

Wickham H, François R, Henry L, Müller K, Vaughan D (2023). _dplyr: A Grammar of Data Manipulation_. R package version 1.1.4, <https://CRAN.R-project.org/package=dplyr>.

Wickham H, Vaughan D, Girlich M (2024). tidyr: Tidy Messy Data. R package version 1.3.1, <https://CRAN.R-project.org/package=tidyr>.

Wilke C (2024). _cowplot: Streamlined Plot Theme and Plot Annotations for ‘ggplot2’_. R package version 1.1.3, <https://CRAN.R-project.org/package=cowplot>.

Woolway, R., Jennings, E., Shatwell, T., Golub, M., Pierson, D., & Maberly, S. (2021). Lake heatwaves under climate change. Nature, 589, 402–407. 10.1038/s41586-020-03119-1.

Yang, H., Hu, Z., Shang, L., Deng, Y., & Tang, Y. (2020). A strain of the toxic dinoflagellate Karlodinium veneficum isolated from the East China Sea is an omnivorous phagotroph. Harmful algae, 93, 101775. 10.1016/j.hal.2020.101775.

